# Sex and size matter: foraging ecology of offshore harbour porpoises in waters around Greenland

**DOI:** 10.1101/2022.03.16.484591

**Authors:** Marie Louis, Jennifer Routledge, Mads Peter Heide-Jørgensen, Paul Szpak, Eline Lorenzen

**Affiliations:** Globe Institute, University of Copenhagen, Denmark; Greenland Institute of Natural Resources, Nuuk, Greenland - Copenhagen, Denmark; Trent University, Peterborough, Ontario, Canada

**Keywords:** stable isotopes, δ^13^C, δ^15^N, cetaceans

## Abstract

Individuals of different sex or age can vary in their resource use due to differences in behaviour, life history, energetic need, or size. Harbour porpoises are small cetaceans that rely on a constant prey supply to survive. Here, we use bone collagen carbon (δ^13^C) and nitrogen (δ^15^N) isotope compositions to elucidate sex and size differences in the foraging ecology of harbour porpoises from West Greenland. In this region, populations have a unique offshore, deep-water ecology. Female harbour porpoises are larger than males and we find that females have a higher trophic level than males, and δ^15^N positively correlates with size for females only. This indicates that size may matter in the ability of females to handle larger prey and/or dive deeper to catch higher trophic level prey. These results suggest that females, which also feed their calves, may be under different ecological constraints than males. We also analysed the harbour porpoise data with comparable stable isotope data from Greenland populations of belugas and narwhals. Consistent with their small body size, and a diet consisting primarily of capelin, we find that harbour porpoises have a lower trophic level than belugas and narwhals. Furthermore, harbour porpoises have the largest ecological niche of the three species, which is in accordance with tagging studies indicating they have a wide range in shelf and deep offshore waters of the sub-arctic and North Atlantic.

## Introduction

Intraspecific variation in foraging ecology may be driven by the distribution and abundance of prey and of interspecific competition, and by intraspecific sex, age, and size differences. Resource partitioning between individuals of different sex or age may reflect differing behaviour, life histories, physiological requirements, or sexual size dimorphism (Hutchinson 1957; Schoener 1974; Bolnick et al. 2003; Bjorkland et al. 2015).

Strong differences in the ecology of females and males, as well as between individuals of different age, have been observed in many pinniped species (Bailleul et al. 2010; Bjorkland et al. 2015; Kernaléguen et al. 2016; Lima et al. 2019). In cetaceans, ontogenic changes in diet may be common (Newsome et al. 2009; Riccialdelli and Goodall 2015; Giménez et al. 2017), yet marked sex differences have mainly been reported in species with relatively strong sexual size dimorphism (Reisinger et al. 2016; Pirotta et al. 2020; Louis et al. 2021). These intraspecific variations in marine mammal ecology have been linked to habitat segregation, maternal investment, sex and age related size differences, or some combination of these. In many species of pinnipeds and toothed cetaceans, males are larger than females (Mesnick and Ralls 2018). Larger individuals may have greater diving abilities and can capture and handle bigger prey, which usually feed at higher trophic levels (Costa et al. 2004).

In some toothed whale species, females are the larger sex (Mesnick and Ralls 2018). For example, in harbour porpoises (*Phocoena phocoena*; small cetaceans inhabiting temperate-to-cold waters in the Northern hemisphere, from the Equator to the Arctic) females are larger and heavier (153-163 cm and 55-65 kg) than males (141-149 cm and 46-51 kg) (Lockyer 2003). Harbour porpoises have high energetic requirements, as they have limited capacity to store energy due a high body surface area:volume ratio. To maintain basal metabolism, they rely on finding prey and eating at predictable and short intervals (Koopman 1998; Wisniewska et al. 2016). They are thus found in productive areas with abundant prey resources.

Feeding studies based on stomach contents in the North Atlantic have shown harbour porpoises are opportunistic, eating mainly fish but also cephalopods and crustaceans (Santos and Pierce 2003), although their prey may be chosen based on calorific value (Spitz et al. 2012). As they are generalist feeders, their diet may reflect prey availability in an ecosystem (Christensen and Richardson 2008). Their feeding ecology has also been studied using stable isotope analyses of δ^13^C and δ^15^N in muscle and/or bone; δ^13^C is a proxy for feeding habitat (pelagic *versus* benthic, offshore *versus* coastal) and δ^15^N is a proxy for trophic level (Peterson and Fry 1987; Hobson et al. 1995; France et al. 1998; Newsome et al. 2010). Studies from several regions in European waters and the Black Sea have shown sex and/or size variation in δ^13^C and/or δ^15^N, but no consistent patterns have been observed across different regions (Das et al. 2003, 2004; Jansen et al. 2012). Differences between sexes have been attributed to (i) seasonal habitat segregation, with females distributed in shallower waters than males, in particular when nursing and calving, (ii) differences in the prey species eaten, or (iii) the size of the prey consumed (Das et al. 2003, 2004; Jansen et al. 2012).

Harbour porpoises are mainly found in continental shelf waters less than 200 m deep. However, recent satellite tracking in West Greenland showed that the species leaves the continental shelf in the autumn and undertakes large-scale movements (up to 2,000 km) in >2,500 m deep, temperate, offshore waters in the North Atlantic in the winter/spring. Most animals return to the continental shelf in West Greenland the next summer while some move to East Greenland (Nielsen et al. 2018, 2019). The genetically and morphologically distinct Greenland population has recently been described as a different “offshore” ecotype, as elsewhere in the northern hemisphere harbour porpoises may be considered more “coastal” (Galatius and Gol’din 2011; Olsen et al. 2022). When offshore, the Greenland harbour porpoises may feed in the mesopelagic layer (Nielsen et al. 2019). When the porpoises are on the shelf of West Greenland their diet consists mainly of capelin (*Mallotus villosus*), although a more diverse diet including Atlantic cod (*Gadus morhua*) and Greenland cod (*G. ogac*) has been observed in stomach contents since 2009 (Lockyer et al. 2003; Heide-Jørgensen et al. 2011).

Here, to elucidate the long-term foraging ecology of the Greenland harbour porpoises, we analysed bone collagen carbon (δ^13^C) and nitrogen (δ^15^N) isotope compositions from individuals collected in Maniitsoq, West Greenland. Specifically, we aimed to investigate: (i) sex-related differences in foraging ecology; (ii) size-related dietary differences, which could reflect variation in prey handling or diving abilities of smaller and larger individuals, (iii) dietary niche of harbour porpoises relative to co-occurring populations of other cetacean species in Greenland, from where comparable bone collagen stable isotope data are available.

## Materials and Methods

### Laboratory analyses

We sampled 300 mg of bone powder from the skulls of 27 sub-adult or adult harbour porpoises for stable isotope analyses of δ^13^C and δ^15^N (Table S1). The specimens were collected during subsistence hunting in Maniitsoq in West Greenland in 1989; voucher specimens (skulls) of all individuals are housed at the Natural History Museum of Denmark, University of Copenhagen. Sex has previously been detemined by inspection of reproductive organs (Kinze 1990; Lockyer et al. 2001); the samples comprised 13 females and 13 males, sex information was not available for one individual.

Powdered samples were lipid extracted with chloroform:methanol and demineralised in 0.5 M HCl, following the procedure described in the supplementary text. We determined carbon and nitrogen stable isotopic and elemental compositions using a Euro EA 3000 Elemental Analyzer (Euro Vector SpA) coupled to a Nu Horizon (Nu Instruments, UK) continuous flow isotope ratio mass spectrometer at the Water Quality Centre at Trent University, Canada. The calibration procedure is described in the supplementary text. We adjusted the δ^13^C values to correct for the change in atmospheric and oceanic dissolved inorganic carbon that has occurred since the late 19th century due to industrialization (the “Suess Effect”; [18,19]), following Szpak et al. 2020 [20], but we used 0.014 for the annual rate at which δ^13^C has declined for a particular water body [21].

### Statistical analyses

All statistical analyses were performed in R v.3.6.1 (R Core Team 2019).

#### Within species

We tested for dietary differences between female and male harbour porpoises (which we term groups) by comparing δ^13^C and δ^15^N using Student’s t-tests, the data satisfied normality and homogeneity of variances.

We used the isotopic niche as a proxy for ecological niche (Bearhop et al. 2004a; Newsome et al. 2007; Jackson et al. 2011), and compared isotopic niches between groups using Bayesian multivariate ellipse-based metrics implemented in the packages SIBER and rjags (Bearhop et al. 2004b; Newsome et al. 2007; Jackson et al. 2011; Plummer and Others 2016). We calculated standard ellipse areas corrected for sample size (SEA_C_), and Bayesian standard ellipses (SEA_B_) for each group. We estimated SEA_B_ using 10^5^ posterior draws, a burn-in of 10^3^ and a thinning of 10, and used SEA_B_ to test for differences in niche width among groups (i.e., the proportion (p) of draws of the posterior distribution of the SEA_B_ in which the area of one group was smaller than the other). We set the prediction ellipses to contain approximately 40% of the data. We evaluated isotopic niche similarity between two groups as the proportion (%) of the non-overlapping area of the maximum likelihood (ML) fitted ellipses of the two.

We investigated the association between the size of individual harbour porpoises and their isotopic composition. For this, we ran a linear regression between skull length, which we considered a proxy for body length in cetaceans (Westgate 2007), and δ^13^C or δ^15^N for all individuals, and separately for females and males. Skull length was measured using a scale, and may be prone to errors given the small size of the skulls. We could not measure two of the individuals because their skulls were fragmented, and hence sample size for this analysis was 25.

#### Among species

To investigate the foraging ecology of harbour porpoises from West Greenland relative to co-occurring cetacean species from which comparative bone-collagen stable isotope data are available, we compared our data with available bone collagen δ^13^C and δ^15^Nvalues from belugas (n = 27) and narwhals (n = 40) collected in West Greenland (data from (Skovrind et al. 2019; Louis et al. 2021)). Our analysis also included data from 39 narwhals from East Greenland, as some satellite tagged harbour porpoises have been shown to move between West and East Greenland (Nielsen et al. 2018). All samples were collected between 1990 and 2007. To ensure that the years of sampling were comparable to the time period where the harbour porpoises were sampled, we also ran the analyses with a reduced sample set of only the individuals sampled between 1990 and 1995 (belugas n = 27, WG narwhals n = 10, EG narwhals n =41). Bone collagen δ13C and δ15N reflect the diet of an individual over multiple years (Wild et al. 2000; Hedges et al. 2007)

We tested for dietary differences among the four populations (porpoises, belugas, WG narwhals, EG narwhals) by comparing their δ^13^C and δ^15^N using a Kruskal-Wallis and a Dunn’s post-hoc tests using the R package dunn.test (Dinno 2017). Our data satisfied normality but not homogeneity of variance for this subdivision. We compared isotopic niches (size differences and overlap) among populations using Standard Ellipse Areas statistics, as described in the previous section for groups (female and male harbour porpoises).

We attempted to incorporate prey data in our analyses (Hansen et al. 2012; Watt et al. 2013; Watt and Ferguson 2015), but the resulting mixing polygon was inconsistent with the isotopic compositions observed in the three cetacean species, after correcting for the Suess effect (Szpak et al. 2020) and for trophic enrichment (Szpak et al. 2012). This indicates the available prey isotopic data may not be representative of what would have been eaten by the consumers due to variation in isotopic values in different tissues, differences in the years of sample collection, differences in isotopic compositions for prey in different regions or habitat types for predators that can forage in different locations, and/or inaccurate isotopic compositions or missing prey sources. In addition, prey data were only available from soft tissues (muscle and skin), which inform on seasonal diet and short term variation in resource use (Browning et al. 2014; Giménez et al. 2016), whereas data retrieved from bone reflect the consumer diet over several years, and inform about long-term differences in diet (Wild et al. 2000; Hedges et al. 2007). As prey data are consistently sampled from some seasons but not from others, this can pose problems for comparisons with consumer data from bone.

## Results

### Within species

Bone collagen δ^13^C did not differ significantly between female and male harbour porpoises (t = 0.13, df = 20.5, P = 0.90, Figure 1a, Table S2). Bone collagen δ^15^N was significantly higher in females than in males (t = 2.35, df = 24, P = 0.03). Although not statistically significant, females had a larger niche (SEA_B_ = 1.05 ‰^2^) than males (SEA_B_ = 0.92 ‰^2^); proportion = 0.91 of the ellipses were larger in females than in males (Figure S1, Table S2). The niche overlap between sexes was 20% (Figure 1a).

**Figure 1.**
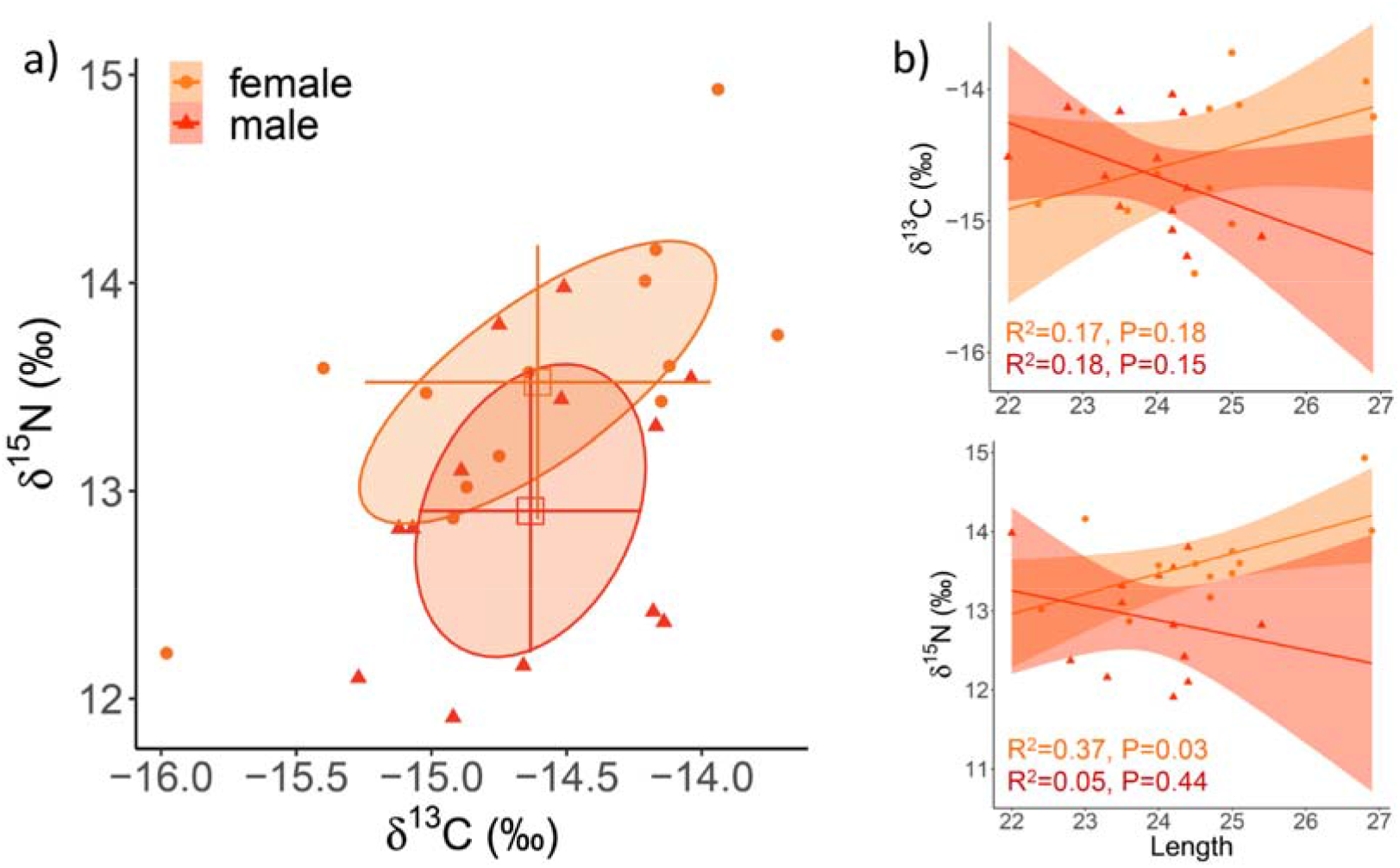
Bone collagen δ^13^C and δ^15^N for 13 female (orange) and 13 male (red) harbour porpoises. In a), solid circles indicate standard ellipse areas encompassing 40% of the data. Mean (square) and SD (error bars) are indicated. b) Variation in δ^13^C and δ^15^N relative to skull length (cm) in either sex. The regression lines for each sex and their confidence intervals (shading) are shown.

Mean skull length was higher in females (m = 24.6 cm, SD = 1.3 cm) than in males (m = 23.9 cm, SD = 0.9 cm), but the differences were not statistically significant (t = 1.72, df = 18.5, P = 0.10). We found no significant correlation between skull length and either δ^13^C or δ^15^N in harbour porpoises (R^2^ = 0.12, P = 0.45 for δ^13^C, R^2^ = 0.12, P = 0.09 for δ^15^N, Figure S2). We did not find a significant correlation when dividing the dataset by sex for δ^13^C (R^2^ = 0.17, P = 0.18 for females, R^2^ = 0.18, P = 0.15 for males, Figure 1b). For δ^15^N, the correlation was significantly positive for females (R^2^ = 0.37, P = 0.03), but not for males (R^2^ = 0.05, P = 0.44, Figure 1b).

### Among species

In the interspecific comparison of four populations, we found significant differences in bone collagen δ^13^C and δ^15^N among populations (n = 133, Kruskal-Wallis X^2^ = 93.8, df = 3, P < 0.01 for δ13C; Kruskal-Wallis X^2^ = 101.0, df = 3, P < 0.01 for δ^15^N, Figure 2b, Table S2).

**Figure 2.**
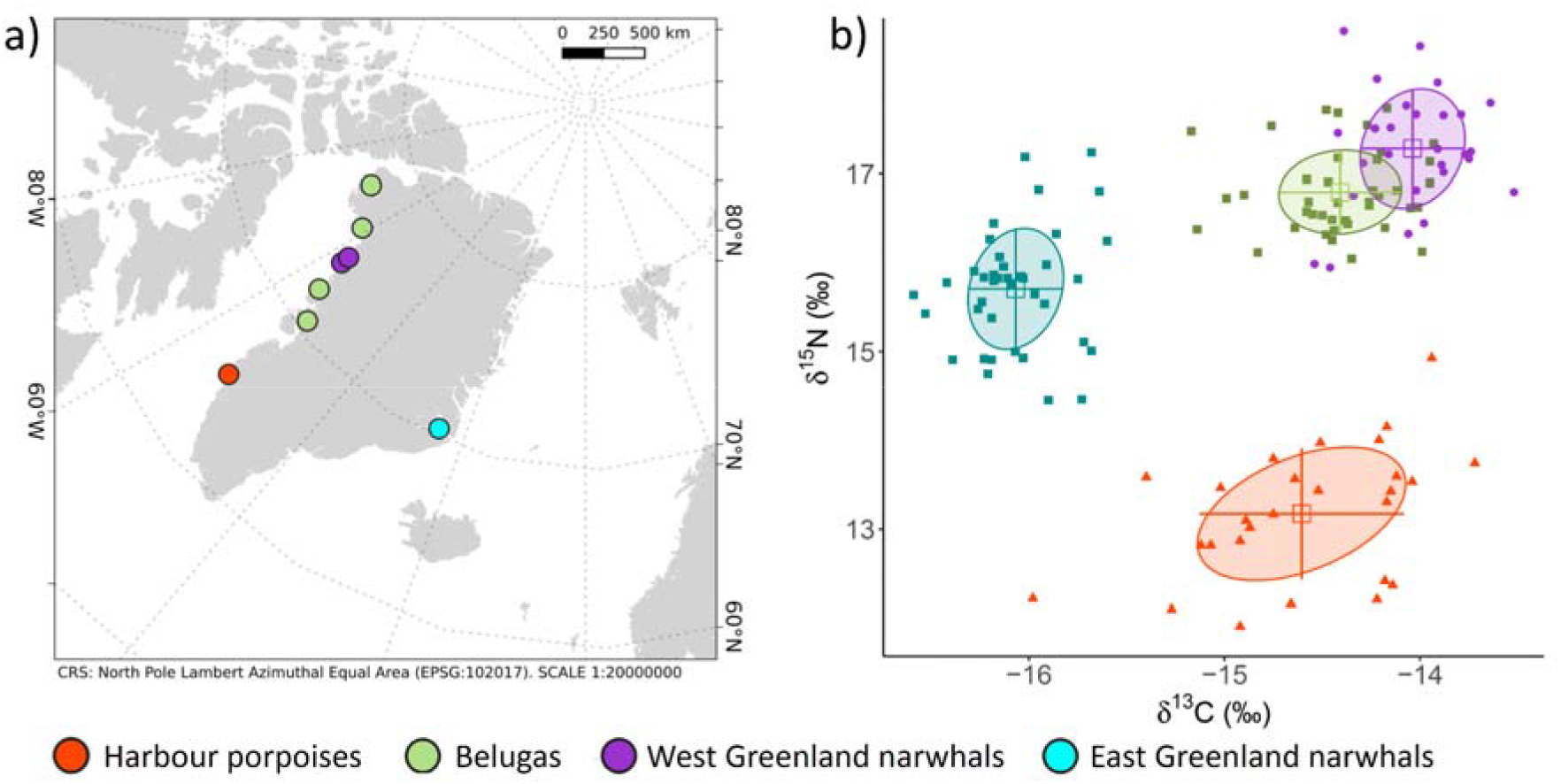
Interspecific comparison of bone collagen δ^13^C and δ^15^N in harbour porpoise, beluga, and narwhal. a) Sample localities, and (b) δ^13^C and δ^15^N values for harbour porpoises (n=27), belugas (n=27), West Greenland narwhals (n=40), and East Greenland narwhals (n=39). In b), solid circles indicate standard ellipse areas encompassing 40% of the data. Mean (square) and SD (error bars) are indicated.

δ^13^C values in harbour porpoises were significantly lower than in belugas and similar to WG narwhals (Dunn test, P = 1.0), and significantly higher than EG narwhals (Dunn test, P < 0.01). δ^15^N values in harbour porpoises were significantly lower than corresponding values in both belugas and narwhals (Dunn test, P < 0.01). The ecological niche of harbour porpoises does not overlap with the other populations, and harbour porpoises have a significantly larger ecological niche than beluga and both narwhal populations (p > 0.99, Figures 2b and S3, Table S2). We find similar results when considering the beluga and narwhal data from the 1990-1995 period only (Figure S4).

## Discussion

We present the first insights into the long-term foraging ecology of harbour porpoises from West Greenland. In contrast with harbour porpoises elsewhere in the northern hemisphere that are year-round inhabitants of continental shelf areas, the porpoises from Greenland represent a population that occur offshore in deep waters for most of the year but spend the summer breeding season on the continental shelf off West Greenland (Nielsen et al. 2018, 2019).

### Within species

Female harbour porpoises are slightly larger than males (Lockyer et al. 2001), and the deep-water habitat of the species in Greenland may strengthen dietary variation of individuals of different sizes, due to differing diving capabilities. We tested for ecological differences between females and males and females have a higher trophic level (δ^15^N) (Figure 1a, Table S2). This finding may reflect their larger size; females may be able to handle larger, higher trophic level prey, or dive deeper and catch higher trophic level prey. In a recent satellite tagging study of harbour porpoises around Maniitsoq, the two (of 19) individuals with the highest median of the daily maximum dive depth on the continental shelf were the largest females (Nielsen et al. 2018). This supports that size may matter in diving ability, and in accessing higher trophic level prey in deeper waters. In belugas and narwhals, which are both found year-round in Arctic waters and also live in deep waters in particular in the winter, males are the larger sex, and bone collagen stable isotope data show that males have a higher trophic level than females (Figure S5) (Louis et al. 2021). This supports our hypothesis that a larger size influences foraging ecology in cetaceans, and may enable greater diving or bigger prey handling capabilities.

Within harbour porpoises, sex differences in foraging ecology have been observed in other geographic regions using stable isotope analysis of soft tissues. For example, in the Black Sea, females have higher muscle δ^13^C than males, possibly due to habitat segregation, as they occur in shallow coastal waters during calving and nursing periods, while males may feed on prey further offshore (Das et al. 2004). In the North Sea, female muscle tissues are enriched in δ^15^N relative to males, and this has also been attributed to a more coastal diet in females relative to males (Das et al. 2003; Jansen et al. 2012). In the Baltic and Kattegat/Skagerrak Seas, and in the Dutch coast of the North Sea, no sex differences in bone collagen stable isotope compositions have been observed in stranded or bycaught animals (Angerbjörn et al. 2006; Jansen et al. 2012). In the southern North Sea, sex differences have been reported based on bone samples collected between 1987 and 2009, with females showing lower δ^15^N than males (Christensen and Richardson 2008). Together with these previous findings, our results indicate that sex variation in diet is population specific, and likely the product of varying ecology, inter- and intrapsecific competition, and prey distribution. For example, porpoises in Greenland have an offshore distribution; 30 tagged individuals around Maniitsoq had a range 8 times bigger than 71 individuals tagged in Denmark (Nielsen et al. 2018). Harbour porpoises in Greenland also showed they can dive to nearly twice the maximum depth previously reported for the species (Nielsen et al. 2018), and are specialised in feeding in the mesopelagic layer (Nielsen et al. 2019). These differences in habitat use and behaviour may explain variation in intraspecific resource partitioning among different populations facing various ecological constraints.

Although not significant (91% of the ellipses are larger in females than in males), we do observe a trend that females have a relatively larger ecological niche than males (Figures 1a and S1). This could reflect that females have higher energetic needs than males as they feed their young for a period of at least 8 months (Lockyer 2003), and hence may need to feed on whatever prey is available and catchable. Harbour porpoises need to feed constantly to maintain basal metabolism (Koopman 1998; Wisniewska et al. 2016). Females with dependent calves have even higher energetic requirements to maintain both their basal metabolism, and to be able to feed their calves.

The cold waters around Greenland may constrain the ecology of harbour porpoises further, and individuals in this region are among the shortest and heaviest (Lockyer 2003). We find no correlation between skull length and foraging habitat (δ^13^C) or trophic level (δ^15^N) (Figure S2). When separating the data by sex, we find a positive and significant correlation between trophic level and size in females only (Figure 1b). This size variation in diet further supports our hypothesis that larger size enables females to catch prey at higher trophic level, due to either higher diving or prey handling capabilities, or both. It is further supported by similar patterns in belugas in West Greenland and in narwhals in East Greenland, where the largest sex only (males are larger than females in both species), has a significant positive correlation between size and trophic level (Louis et al. 2021). In studies from other regions, including the North Sea, where harbour porpoises have a coastal and shallow-water distribution, the pattern is reversed, i.e. larger individuals have a lower trophic level (Jansen et al 2012). This indicates that size variation in foraging ecology is also population specific, and that size matters differently for populations with different ecologies such as ranging in deep or shallow waters.

### Among species

To investigate resource partitioning among cetaceans in the waters around Greenland, we compared the ecological niche of harbour porpoises with that of belugas and narwhals, from which comparable bone collagen stable isotope data are available (Louis et al. 2021). Belugas and narwhals (Monodontidae) are medium-sized whales that inhabit the Arctic year-round. The most southern distributions of the two species in the Baffin Bay area, and of narwhals in East Greenland (where beluga are not found), overlap spatially but not temporally with harbour porpoises (Nielsen et al. 2018; Hobbs et al. 2019). While harbour porpoises are found in the Baffin Bay area in summer, belugas and narwhals utilise the area in winter. Both belugas and narwhals show extreme site fidelity to migration routes between coastal summering and offshore deep water wintering grounds. Some prey of belugas and narwhals, such as capelin and Arctic cod (*Boreogadus saida*), have also been identified as important prey for harbour porpoises in the area (Heide-Jørgensen and Teilmann 1994; Lockyer et al. 2003; Heide-Jørgensen et al. 2011; Watt et al. 2013). Harbour porpoises in Greenland show site fidelity but to a lesser extent than belugas and narwhals. Of six tagged individuals with data spanning more than a year, all returned to the tagging area around Maniitsoq. Four individuals did so the summer after capture, while two males moved to either East Greenland or the North Atlantic, and returned two years later (Nielsen et al. 2018).

We find that harbour porpoises occupy a distinct ecological niche (Figure 1b). The trophic level of harbour porpoises, represented by δ^15^N, is much lower than those of belugas and narwhals (Figure 2, Table S2). This finding is in agreement with harbour porpoises caught around Maniitsoq and surrounding regions between 1987-1995 (our samples are from 1989) having a diet mainly composed of capelin (Lockyer et al. 2003; Heide-Jørgensen et al. 2011). Belugas and narwhals both feed on capelin, but also on larger prey at higher trophic level, including Arctic cod (*Boreogadus saida*), polar cod (*Arctogadus glacialis*) and halibut (*Reinhardtius hippoglossoides*) (Heide-Jørgensen and Teilmann 1994; Laidre and Heide-Jorgensen 2005; Marcoux et al. 2012; Watt et al. 2013). Furthermore, belugas and narwhals are approximately three times the size of a harbour porpoise, and are able to catch and handle much larger prey, which are usually at a higher trophic level.

δ^13^C values in harbour porpoises are significantly lower than in belugas, and are similar to West Greenland narwhals (Figure 2, Table S2). This pattern suggests that harbour porpoises, akin to narwhals, have a more pelagic diet than belugas, or that they feed further offshore. Our findings are supported by tagging of harbour porpoises caught in Maniitsoq, which show they feed in the mesopelagic layer (Nielsen et al. 2019). The range of the δ^13^C values in harbour porpoise individuals is large (min:-15.98, max:-13.72). The lowest observed δ^13^C values are similar to those in East Greenland narwhals. We hypothesised that the variation in δ^13^C values found in narwhals between West and East Greenland may reflect regional variation at the base of the food web (Louis et al. 2021). Thus, low δ^13^C values for some harbour porpoises may reflect that some individuals caught in Maniitsoq in West Greenland moved to East Greenland (Nielsen et al. 2018). We were unable to include prey isotopic data in our analyses to further investigate the dietary niche of the three cetacean species, as only data from soft tissues are available. Prey isotope data may vary seasonally for soft tissues, and therefore, if not sampled evenly across all seasons, data may be biased towards one period of time. This presents a problem for comparisons with stable isotope data from the consumers retrieved from bone collagen, which reflect average diet over multiple years (see Material and Methods for further details).

Harbour porpoises also have a wider ecological niche than belugas and narwhals (Figures 2 and S3, Table S2), despite all 27 harbour porpoise specimens being sampled from one location (Maniitsoq), and collected within a single year (1989). The beluga and narwhal samples spanned either several locations, several years, or both. The beluga samples were collected between 1990 and 1994. The narwhal samples were collected during two time periods: 1993-1995 and 2002-2007, with about half the samples collected during each period. Restricting the narwhal data to the 1993-1995 period does not affect our findings (Figure S4).

The results are consistent with tracking studies indicating that although harbour porpoises tagged around Maniitsoq in West Greenland show site fidelity, they have a large range (i.e. the minimum convex polygons from 30 animals was around 4.2 million km^2^) in the North Atlantic and can move between West and East Greenland in offshore and deep waters (Nielsen et al. 2018). Additionally, different individual harbour porpoises show varying movement patterns (Nielsen et al. 2018). Although belugas and narwhals also make large-scale movements, they show very strong site fidelity to the same migratory routes they follow year after year (Hobbs et al. 2019). These habitat use and behavioural differences likely explain why the isotopic niche of harbour porpoises is larger than those of the Monodontids. In addition, unlike harbour porpoises, no movements of narwhals have been so far recorded between West and East Greenland (Heide-Jørgensen et al. 2013, 2015).

## Conclusions

We find sex and size variation in the foraging ecology of harbour porpoises. The females are the larger sex and have a higher trophic level than males. Females also show size variation in δ^15^N. This suggests that size matters in catching higher trophic level prey, and that females are under other ecological constraints than males. Our study shows that harbour porpoises feed a lower trophic level than belugas and narwhals, reflecting ecological differences among species, and differences in the size of consumed prey. Greenland harbour porpoises have a larger isotopic niche than belugas and narwhals, likely due to their wider distributional range in waters of the sub-arctic and North Atlantic, and more individual variation in habitat use (Nielsen et al. 2018). The samples we analysed were collected in 1989; a concurrent study based on stomach contents showed harbour porpoises caught in Maniitsoq, Nuuk, and Paamiut (all in West Greenland) between 1987 and 1995 had a diet primarily consisting of capelin (Lockyer et al. 2003). Due to an increase in prey abundance associated with rising sea temperatures, harbour porpoises in these West Greenland locations have more recently been feeding on a more diverse range of species, including a larger proportion of Atlantic cod and Greenland cod (Heide-Jørgensen et al. 2011). This dietary shift to higher trophic level prey has likely resulted in improved body condition of the porpoises, and may also have affected intraspecies variation in foraging ecology. Analysing bone collagen stable isotopes of individuals collected in more recent years may shed light on how these more recent changes in climate are driving shifts in resource partitioning within harbour porpoises.

## Acknowledgements

We thank Peter Rask Moller and Daniel Klingberg Johansson for access to the harbour porpoise collections of the Natural History Museum of Denmark, University of Copenhagen, and in locating the specimens. We also thank the hunters who collected the specimens in Maniitsoq, Greenland. We thank René Swift for making the map for Figure 1. ML was supported by a postdoctoral fellowship from the Greenland Research Council.

## Author Contributions

ML conceived the study; PS and JR ran the laboratory analyses; ML analysed the data; ML, PS, MPH and EDL interpreted the data; ML and EDL wrote the manuscript, with input from all co-authors. All authors approved the final manuscript.

## Notes

### Competing Interest Statement

The authors have declared no competing interest.

