## Supplementary material for "Sex and size matter: foraging ecology of offshore harbour porpoises in waters around Greenland"

Electronic supplementary material

Supplementary text

**Stable isotope laboratory analyses**

Powdered samples underwent lipid extraction with 3 repetitions of 10 ml 2:1 chloroform/methanol (v/v) under sonication for 1 h. After removing the solvent, we dried the samples under normal atmospheric pressure for 24 h. We demineralized the samples in 10 ml of 0.5 M HCl for 1 h while agitating them by an orbital shaker. After demineralization, we rinsed the samples to neutrality with Type I water, and heated them at 75 °C for 36 h in 10^−3^ M HCl to solubilize the collagen. We freeze dried the water soluble collagen. Upon analysis there was an observed correlation between *δ*^13^C and C:N_atomic_, indicating the presence of residual lipids. To extract these lipids we solubilized the lyophilized collagen and treated with 10 ml 10:5:4 chloroform/methanol/water (v/v/v) under sonication for 1 h. After centrifugation, the layer containing solubilized collagen, and methanol was retained and the methanol was evaporated from the solution at 60 °C for 24 h. We freeze dried and weighed the samples before adding 0.5 mg into tin capsules for elemental and isotopic analysis. We determined carbon and nitrogen stable isotopic and elemental compositions using a Euro EA 3000 Elemental Analyzer (Euro Vector SpA) coupled to a Nu Horizon (Nu Instruments, UK) continuous flow isotope ratio mass spectrometer at the Water Quality Centre at Trent University, Canada. We calibrated stable carbon and nitrogen isotopic compositions relative to the VPDB and AIR scales using USGS40 and USGS41a (glutamic acid, table S3) [[1,2]](https://paperpile.com/c/DpSwmj/FlEa3+2Ibwt). Analytical uncertainty was monitored using four internal reference materials in addition to USGS40 and USGS41a: SRM-1 (caribou bone collagen, long-term average *δ*^13^C = −19.40±0.08 ‰, *δ*^15^N = +1.82±0.11 ‰), SRM-2 (walrus bone collagen, long-term average *δ*^13^C = −14.82±0.06 ‰, *δ*^15^N = +15.59±0.14 ‰), SRM-14 (polar bear bone collagen, long-term average *δ*^13^C = −13.68±0.08 ‰, *δ*^15^N = +21.61±0.16 ‰), and SRM-17 (phenylalanine, long-term average *δ*^13^C = −12.45±0.04 ‰, *δ*^15^N = +3.17±0.15 ‰). The observed isotopic compositions for these standards are presented in table S4. For samples analyzed in duplicate, the mean difference between pairs was 0.06 ‰ for *δ*^13^C and 0.11 ‰ for *δ*^15^N. Standard uncertainty was determined to be ±0.11 ‰ for *δ*^13^C and ±0.21 ‰ for *δ*^15^N. We also adjusted the *δ*^13^C values to correct for the change in atmospheric and oceanic dissolved inorganic carbon that has occurred since the late 19th century due to industrialization (the “Suess Effect”; [[3,4]](https://paperpile.com/c/DpSwmj/x963i+hQQi5)), following [[5]](https://paperpile.com/c/DpSwmj/c7PR5), although we used 0.014 for the annual rate at which δ13C has declined for a particular water body [[6]](https://paperpile.com/c/DpSwmj/be1O).

Supplementary figures and tables

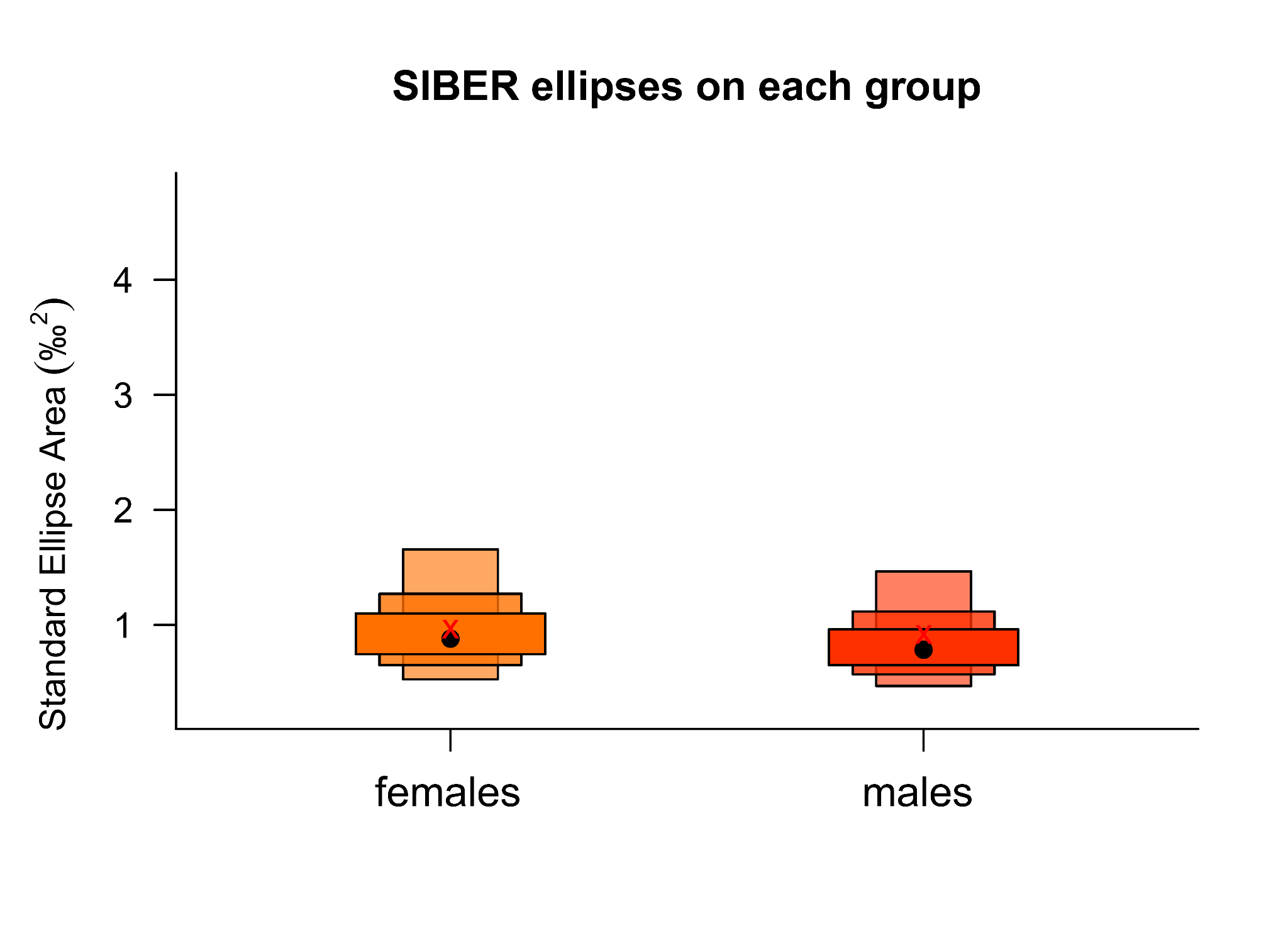

Figure S1. Size of the Standard Ellipse Areas (SEA, ‰^2^) for 13 female and 13 male harbour porpoises collected in Maniitsoq, West Greenland. The black dot indicates the mean size of the Bayesian standard ellipse areas (SEA_B_), the red cross the mean for the standard ellipse areas corrected for sample size (SEA_C_), and the box edges the 50, 75, and 95% credible intervals.

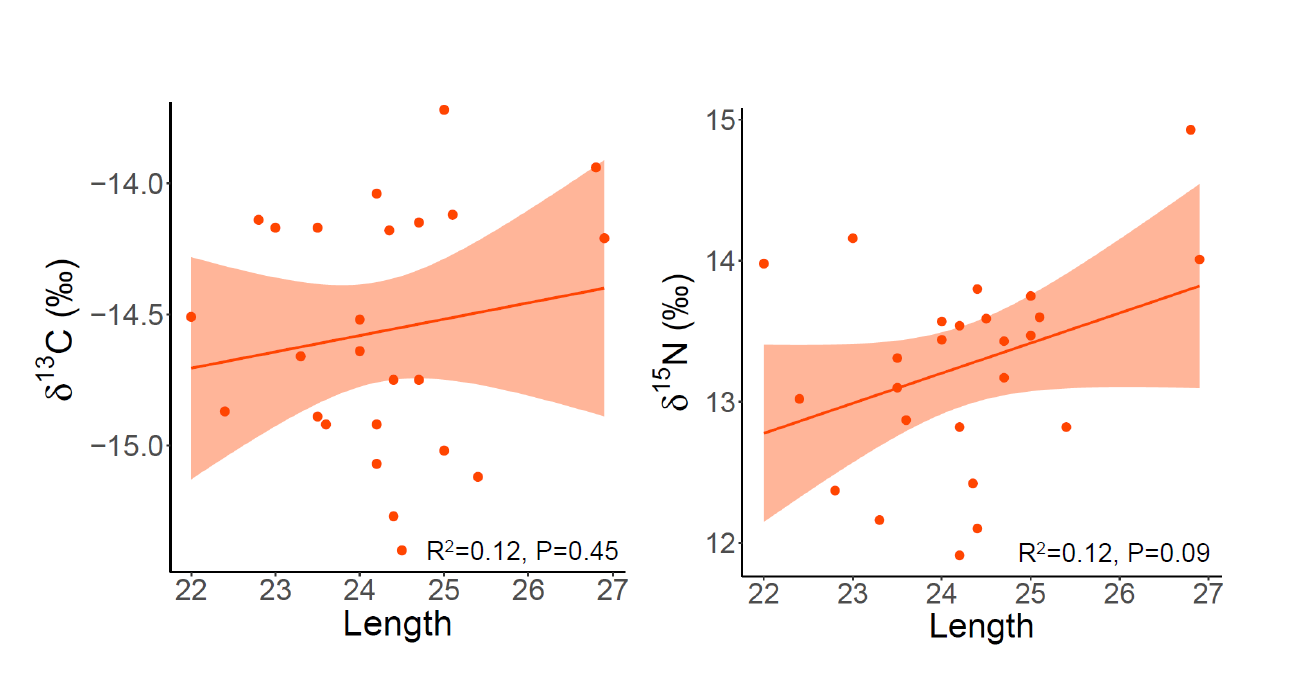

Figure S2. Variation in bone collagen *δ*^13^C and *δ*^15^N of 25 harbour porpoises according to skull length (in cm).

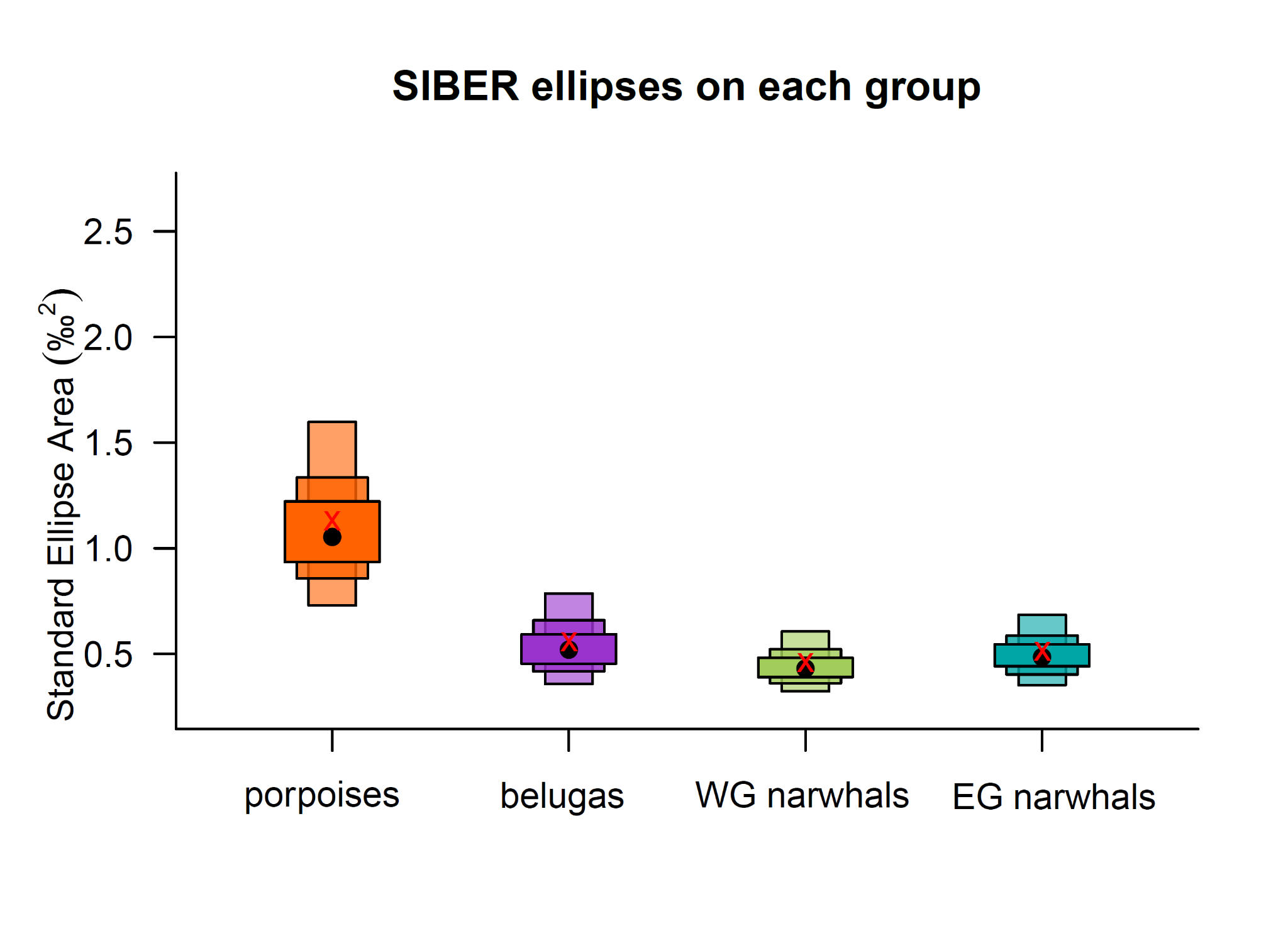

Figure S3. Size of the Standard Ellipse Areas (SEA, ‰^2^) for 27 harbour porpoises, 27 belugas, 40 West Greenland (WG) narwhals, and 39 East Greenland (EG) narwhals. Black black dot indicates mean size of the Bayesian standard ellipse areas (SEA_B_), red cross indicates mean for the standard ellipse areas corrected for sample size (SEA_C_), box edges indicate 50, 75, and 95% credible intervals.

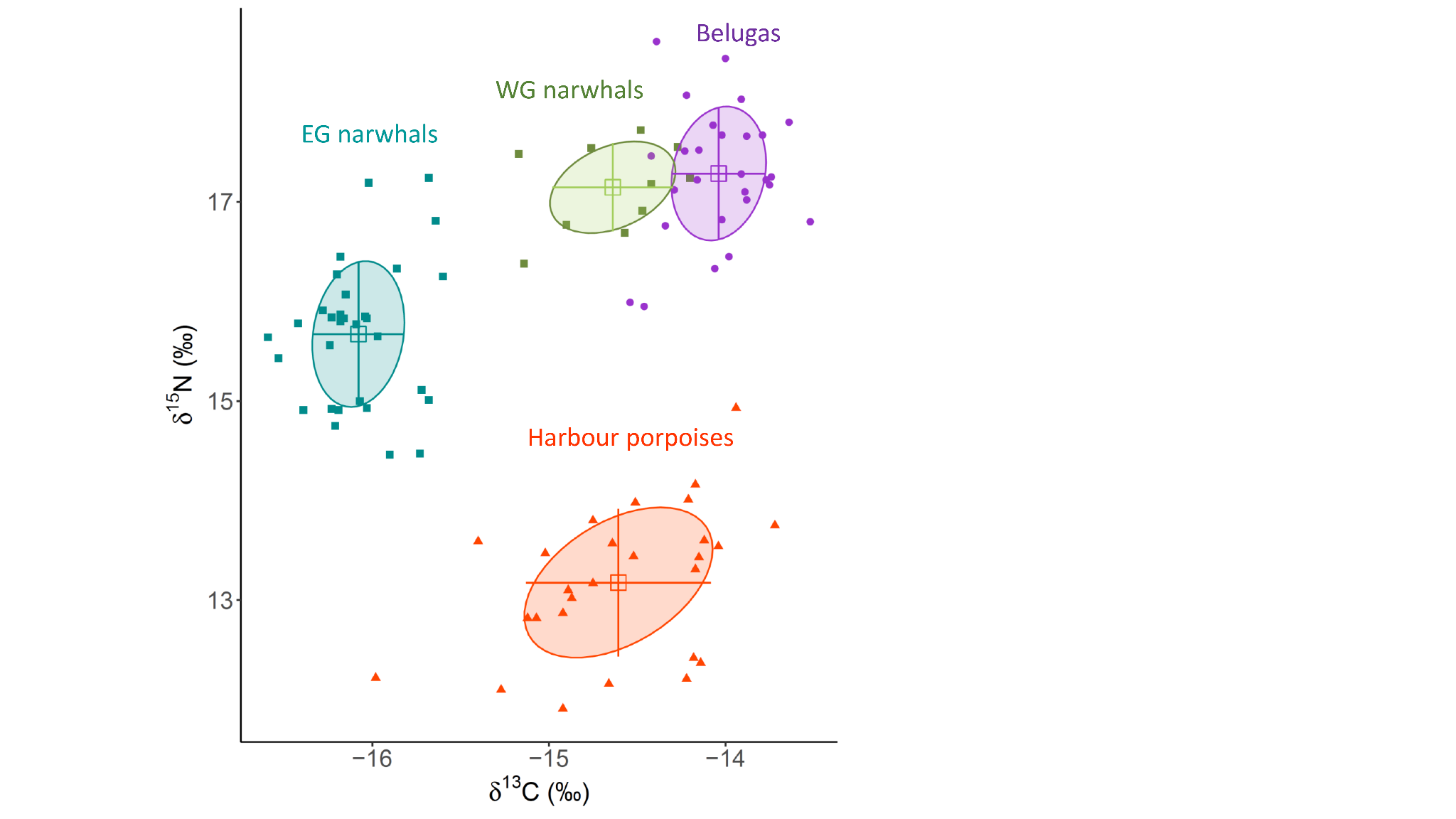

Figure S4. Bone *δ*^13^C and *δ*^15^N for 27 harbour porpoises, 27 belugas, and a reduced dataset of narwhal specimens collected in 1993-1995: 10 West Greenland (WG) narwhals and 31 East Greenland (EG) narwhals. Solid circles indicate standard ellipse areas encompassing 40% of the data. Mean (square) and SD (error bars) are indicated.

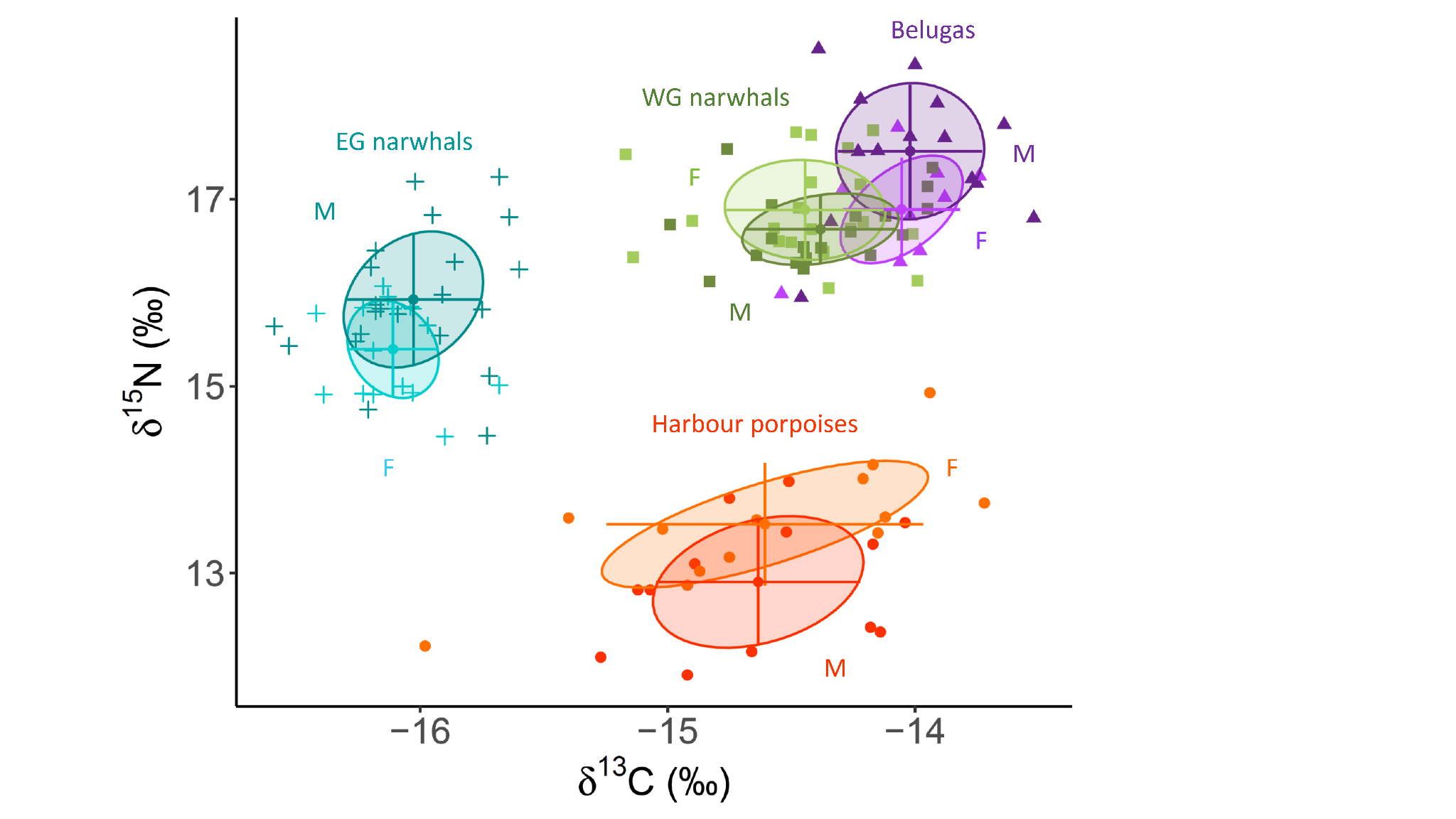

Figure S5. Bone collagen *δ*^13^C and *δ*^15^N for female and male harbour porpoises (n_F_=13, n_M_=13), belugas (n_F_=9, n_M_=14), WG narwhals (n_F_=20, n_M_=19), and EG narwhals (n_F_=16, n_M_=22). Solid circles indicate standard ellipse areas encompassing 40% of the data. Mean (square) and SD (error bars) are indicated.

Table S1. Sample information of harbour porpoise specimens analysed in the study. For each individual, we include, TEAL (laboratory ID at the University of Trent, Canada), sample ID (specimen ID at the Natural History Museum Copenhagen), *δ*^13^C, *δ*^13^C corrected for the Suess effect, *δ*^15^N, C:N Atomic ratio, Sampling year, Sex, and Skull length. The latter is used as a size proxy for each individual (in cm, length of total skull length).

| **TEAL** | **Sample ID** | ***δ^13^C***  (‰) | **δ^13^C Suess** (‰) | ***δ^15^N*** (‰) | **C:N** | **Sampling year** | **Sex** | **Skull length (cm)** |
| --- | --- | --- | --- | --- | --- | --- | --- | --- |
| 13363 | CN618 | -15.24 | -14.64 | 13.57 | 3.15 | 1989 | F | 24 |
| 13362 | CN621 | -15.35 | -14.75 | 13.17 | 3.25 | 1989 | F | 24.7 |
| 13341 | CN625 | -14.76 | -14.17 | 14.16 | 3.17 | 1989 | F | 23 |
| 13340 | CN627 | -15.46 | -14.87 | 13.02 | 3.27 | 1989 | F | 22.4 |
| 13344 | CN628 | -15.11 | -14.51 | 13.98 | 3.28 | 1989 | M | 22 |
| 13343 | CN629 | -14.71 | -14.12 | 13.6 | 3.21 | 1989 | F | 25.1 |
| 13338 | CN630 | -15.48 | -14.89 | 13.1 | 3.19 | 1989 | M | 23.5 |
| 13345 | CN631 | -14.63 | -14.04 | 13.54 | 3.19 | 1989 | M | 24.2 |
| 13342 | CN632 | -15.87 | -15.27 | 12.1 | 3.46 | 1989 | M | 24.4 |
| 13339 | CN633 | -14.82 | -14.22 | 12.21 | 3.12 | 1989 | U | NA |
| 13348 | CN634 | -15.51 | -14.92 | 11.91 | 3.34 | 1989 | M | 24.2 |
| 13351 | CN635 | -15.35 | -14.75 | 13.8 | 3.38 | 1989 | M | 24.4 |
| 13350 | CN636 | -15.12 | -14.52 | 13.44 | 3.24 | 1989 | M | 24 |
| 13352 | CN639 | -14.77 | -14.18 | 12.42 | 3.13 | 1989 | M | 24.35 |
| 13346 | CN641 | -14.32 | -13.72 | 13.75 | 3.14 | 1989 | F | 25 |
| 13347 | CN642 | -14.8 | -14.21 | 14.01 | 3.26 | 1989 | F | 26.9 |
| 13349 | CN643 | -16 | -15.4 | 13.59 | 3.46 | 1989 | F | 24.5 |
| 13359 | CN645 | -15.51 | -14.92 | 12.87 | 3.2 | 1989 | F | 23.6 |
| 13361 | CN649 | -14.73 | -14.14 | 12.37 | 3.12 | 1989 | M | 22.8 |
| 13364 | CN651 | -15.26 | -14.66 | 12.16 | 3.08 | 1989 | M | 23.3 |
| 13357 | CN652 | -16.57 | -15.98 | 12.22 | 3.35 | 1989 | F | NA |
| 13358 | CN655 | -15.66 | -15.07 | 12.82 | 3.29 | 1989 | M | 24.2 |
| 13360 | CN656 | -14.76 | -14.17 | 13.31 | 3.22 | 1989 | M | 23.5 |
| 13355 | CN657 | -15.72 | -15.12 | 12.82 | 3.33 | 1989 | M | 25.4 |
| 13353 | CN658 | -15.61 | -15.02 | 13.47 | 3.31 | 1989 | F | 25 |
| 13354 | CN660 | -14.54 | -13.94 | 14.93 | 3.12 | 1989 | F | 26.8 |
| 13356 | CN661 | -14.74 | -14.15 | 13.43 | 3.15 | 1989 | F | 24.7 |

Table S2. Stable isotope data summary. Sample size (n), *δ*^13^C and *δ*^15^N mean and SD (‰), mean standard ellipse areas corrected for small sample size (SEA_C_) and Bayesian standard ellipse areas (SEA_B_) values (‰^2^) in harbour porpoises, belugas, West Greenland (WG) narwhals, and East Greenland (EG) narwhals for all individuals, females and males.

| **Group** | **n** | ***δ*^13^C (‰)** | ***δ*^15^N (‰)** | **SEA_C_ (‰^2^)** | **SEA_B_ (‰^2^)** |
| --- | --- | --- | --- | --- | --- |
| harbour porpoises | 27 | -14.6 (0.52) | 13.2 (0.74) | 1.13 | 1.14 |
| belugas | 27 | -14.0 (0.26) | 17.3 (0.66) | 0.56 | 0.56 |
| WG narwhals | 40 | -14.6 (0.40) | 16.8 (0.46) | 0.46 | 0.46 |
| EG narwhals | 39 | -16.1 (0.24) | 15.7 (0.67) | 0.51 | 0.51 |
| harbour porpoise males | 13 | -14.6 (0.31) | 12.9 (0.68) | 0.92 | 0.92 |
| harbour porpoise females | 13 | -14.6 (0.64) | 13.5 (0.66) | 0.96 | 1.05 |
| beluga males | 14 | -14.0 (0.29) | 17.5 (0.71) | 0.7 | 0.85 |
| beluga females | 9 | -14.1 (0.24) | 16.9 (0.55) | 0.41 | 0.27 |
| WG narwhal males | 19 | -14.4 (0.31) | 16.7 (0.37) | 0.37 | 0.37 |
| WG narwhal females | 20 | -14.4 (0.32) | 16.9 (0.52) | 0.55 | 0.55 |
| EG narwhal males | 22 | -16.0 (0.28) | 15.9 (0.71) | 0.62 | 0.63 |
| EG narwhal females | 16 | -16.1 (0.18) | 15.4 (0.51) | 0.3 | 0.3 |

Table S3. Standard reference materials used for calibration of  *δ*^13^C relative to VPDB and  *δ*^15^N relative to AIR.

| Standard | Material | Accepted *δ*^13^C  (‰, VPDB) | Accepted *δ*^15^N  (‰, AIR) |
| --- | --- | --- | --- |
| USGS40 | Glutamic Acid | −26.39 ± 0.04 | −4.52 ± 0.06 |
| USGS41a | Glutamic Acid | +36.55 ± 0.08 | +47.55 ± 0.15 |

Table S4. Standard reference materials used to monitor internal accuracy and precision.

| Standard | Material | Mean *δ*^13^C  (‰, VPDB) | Mean *δ*^15^N  (‰, AIR) |
| --- | --- | --- | --- |
| SRM−1 | Caribou bone collagen | −19.36 ± 0.11 | +1.81 ± 0.10 |
| SRM−2 | Walrus bone collagen | −14.76 ± 0.12 | +15.59 ± 0.11 |
| SRM−4 | Gluten | −26.76 ± 0.08 | +5.25 ± 0.11 |
| SRM−14 | Polar bear bone collagen | −13.66 ± 0.06 | +21.63 ± 0.11 |
